# Red fluorescent sensor for intensity-and lifetime imaging of cAMP in living cells

**DOI:** 10.64898/2026.11.04.764060

**Authors:** Matea Ivanicic, Lior Tiroshi, Geoffrey Gourinchas, Kengo Adachi, Ryohei Yasuda, Daniel A. Dombeck, Oliver Griesbeck

## Abstract

cAMP (cyclic adenosine monophosphate) is a crucial intracellular messenger involved in neuromodulation and plasticity, but more information on its spatiotemporal regulation is desirable. Here we constructed a red fluorescent indicator based on a truncated cAMP binding site derived from a regulatory PKA subunit of protozoan origin. It contains a single binding site for cAMP highly specific for cAMP over cGMP. chILI (**c**AMP **h**ighly-responsive **I**ntensity and **L**ifetime **I**ndicator) can be imaged as an intensity-based probe, but alternatively also offers options for fluorescence lifetime imaging. When expressed in the mouse brain using AAV-vector mediated gene transfer, it reports spontaneous and evoked cAMP transients from CA1 neurons in the hippocampus of awake mice. chILI will add valuable options for monitoring cAMP in vitro and in vivo.

## Introduction

Cyclic adenosine monophosphate (cAMP)^1^ is a central second messengers relaying the signals of hormones and neurotransmitters into cells. Ever since the identification of the learning mutants *Dunce*^2^ and *Rutabaga*^3^ in Drosophila components of the cAMP signaling pathway have been considered important players in neuronal plasticity across species. The gene underlying the *Dunce* phenotype, for example, codes for a cAMP phosphodiesterase^4^, stressing the importance of localizing and restricting cAMP signals within cells and tissues. Knowledge on spatiotemporal aspects of cAMP signaling in vivo is still rather incomplete, but is expected to increase and profit from advancing imaging and sensing technology. Optical imaging of cAMP started with FRET (Förster Resonance Energy Transfer)-based sensors consisting of the regulatory and catalytic subunits of Protein Kinase A (PKA), labeled with fluorescein and rhodamine, respectively^5–7^. Soon thereafter first GFP-based versions were generated^8^ which abolished the need of covalent labeling of exogeneous PKA subunits and the cumbersome microinjections of sensors into cells of interest. Since, a plethora of cAMP biosensors were developed in various configurations^9^. By now cAMP sensors are starting to be used in vivo for imaging in model organisms^9^. A recent focus lies on developing green^10–14^ and red fluorescent intensity sensors^15–17^ for in vivo work and multiplexing with other types of sensors.

chILI (**c**AMP **h**ighly-responsive **I**ntensity and **L**ifetime **I**ndicator) is a red fluorescent cAMP sensor that can be multiplexed with green activity sensors. To build it, we inserted circularly permuted mApple into a truncated high-affinity cAMP-binding motif of the regulatory subunit of protein kinase A from heterologous *Plasmodium Falciparum*. The resulting indicator has a single cAMP-binding site. The sensor is extremely specific for cAMP over cGMP and was optimized for high brightness and fluorescence change using directed evolution schemes and alternating microbial and mammalian expression and selection systems.

## Results

### Engineering of chILis

To build chILI, we chose the regulatory domain of PKA (Protein Kinase A) from Plasmodium falciparum^18^, a heterologous cAMP binding domain of protozoan origin. Based on insights from its high-resolution crystal structure^19^, pfPKA could be truncated to generate a shortened domain comprising amino acid residues 297-441 that contained a single high affinity cAMP binding site. To generate a red fluorescent sensor, we inserted circularly permuted mApple^20, 21^ into the truncated domain of pfPKA between residues 412 and 413 (original numbering of pfPKA residues). The insertion of cpmApple between positions 412/413 generated a prototypic parental cAMP sensor (chILI.0) with a moderate maximal fluorescence (ΔF/F ∼25% in vitro, and ∼120 % in HEK cells,) (Fig. 1A). Relative brightness of chILi.0 in the bound state was low, with a strong contribution of the neutral chromophore state, suggesting incomplete chromophore transition into the anionic state. We optimized chILIs using various schemes of diversification in screening in microbial and mammalian expression systems. First rounds of directed evolution of chILi.0 were fully performed in mammalian cells. We used a method that relies on CRISPR-Cas9-based introduction of diversity at selected targeted sites^22^ (Fig. S1). We first added a nuclear export signal (NES) to the C-terminus of chILI.0 and fused chILI.0-cNES to mTagBFP2. Then, we deleted 56 nucleotides comprising the 5’ interface between cpmApple and truncated pfPKA (linker 1, L1) to create a shift in the reading frame (Fig. S1A). The resulting non-fluorescent construct was stably integrated in a HEK293FRT Cas9 cell line. To generate a library, cells were co-transfected with an appropriate single guide RNA (sgRNA) and a single strand DNA oligonucleotide (ssODN) carrying four diversified codons sandwiched between homology arms for homology-directed repair (HDR) (Fig. S1A). Incorporation of the library via HDR restored reading frame and generated library cells with red and blue fluorescence (Fig.S1A). Library cells (about 300 000) were collected via fluorescence activated cell sorting (FACS) in high cAMP condition (stimulated with 50 µM forskolin and 50 µM iBMX), initially based on blue fluorescence (Fig. S1B). Cells were allowed to recover from sorting and expanded before being used for a new round of FACS sorting. In the next two rounds we collected cells with higher red-blue ratio in high cAMP condition, with sorting criteria being more restrictive in each round. In the fourth round, dimmer cell population was collected in basal conditions to exclude “always-on” variants. In the last round, cells were measured in high cAMP conditions and single cells from the brightest 1% of the population were sorted into 96-well plates (Fig. S1B). Single variants were then evaluated via automated high-content screening (Fig. S1C) after forskolin and iBMX stimulation. The gene for the highest responding variant chILI.01 was isolated from the genome of the selected cell clone and further characterized (Fig. S1D). chILI.01 had an optimized linker 1 sequence in which the original amino acid sequence GPPV was changed to ECFV. An additional amino acid exchange was identified off-site (F111L). When compared to the parental chILi.0 variant (Fig. S1E), chILI.01 had a dimmer basal fluorescent state and a brighter bound state. This increased the maximal fluorescence change ΔF/F obtained in HEK cells after forskolin/IBMX stimulation from 120% to 440%. Subsequently we attempted to optimize the C-terminal interface region (linker 2, L2) (Fig. 1A) using the identical approach. Any further intervention in this region resulted in loss of fluorescence or worse-performing variants. We reasoned that the engineering of the L1 region already led to an optimized interface environment complementary to L2, with any further changes in the nearby L2 region, being counter-productive. Instead, we chose to add rounds of directed evolution and selection in E. coli using random mutagenesis of the chILI.01 gene. Overall, we next performed 13 rounds of iterative mutagenesis and screening, and screened more than 32000 variants. Based on their basal fluorescence in E. coli, we selected and purified 192 different variants. Selection was further narrowed according to fluorescence response to 1 mM cAMP, extinction coefficient in cAMP-bound state and/or affinity. Of these, we validated 60 promising variants in HEK293T cells. The amino acid exchanges we found beneficial involved I45M/K96M/L111Q/F118L/G199C/E217V/T362P (Fig. S2) and were all accumulated in a variant called chILi.02. Finally, the mutation L232V (Fig. S2) helped to double the brightness of the bound state. This variant was named chILI.1. The development of brightness, fluorescence change and Kds during engineering of chILIs is summarized in Figure 1G.

**Figure 1.**
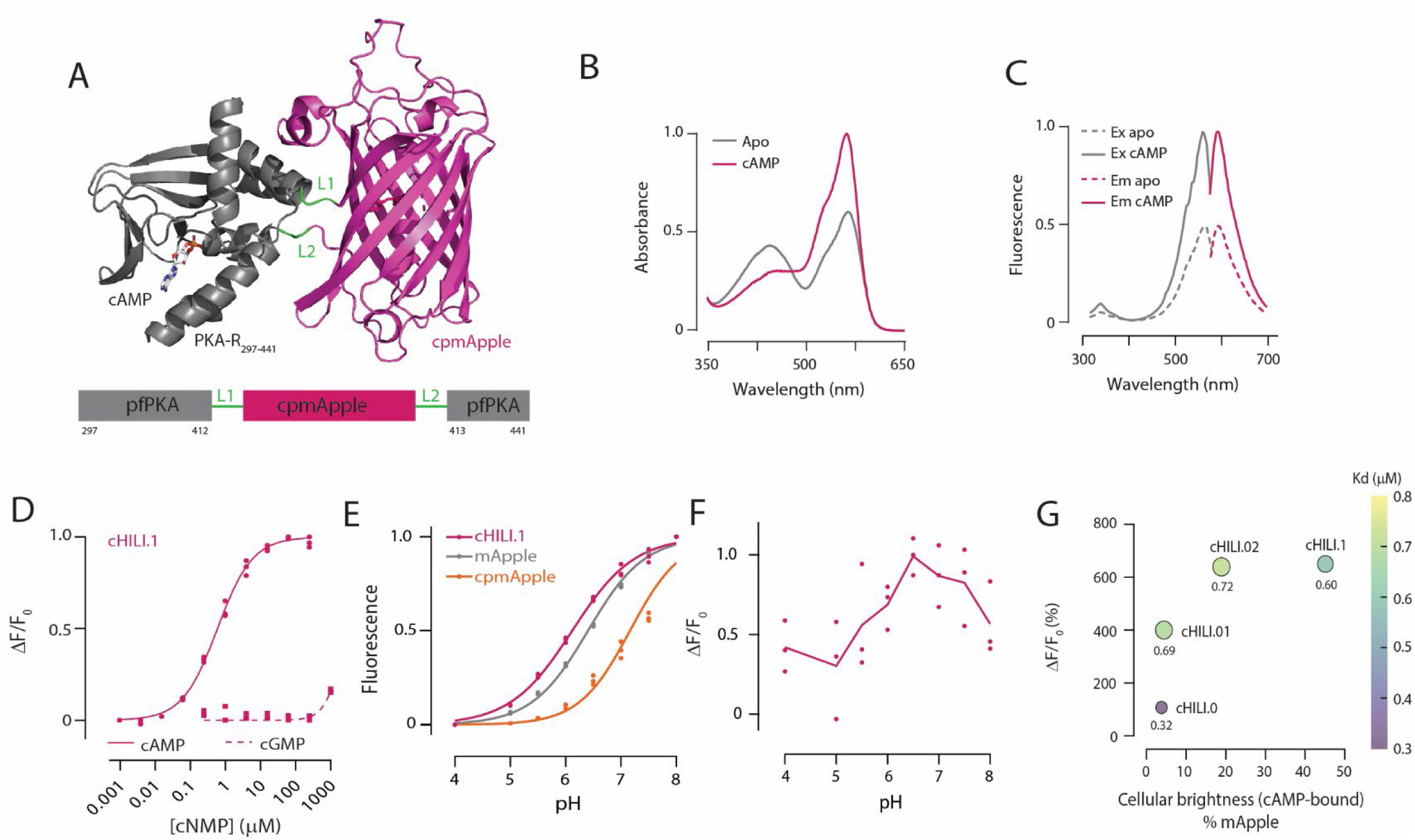
cHILI.1, a red fluorescent cAMP indicator. **A)** AlphaFold3 prediction of the structure of cHILI.1 in the cAMP-bound state (top) and a schematic of the indicator architecture (bottom). cpmApple (magenta), was inserted between amino acid residues 412 and 413 of pfPKA (grey), a truncated regulatory subunit of protein kinase A (PKA) from *Plasmodium falciparum*. L1, L2 (green), linking amino acids between pfPKA and cpmApple. **B)** Absorbance and **C)** excitation and emission spectra of recombinant cHILI.1 in apo and cAMP-bound state. Curves are mean of triplicate protein productions. **D)** Normalized cAMP (dots, full line) and cGMP (squares, dashed line) titration curves of cHILI.1. The Kd of chILI.1 was 600 nM. **E)** Normalized fluorescence emission at different pH in vitro. cHILI.1 was measured in the presence of 1 mM cAMP. **F)** Maximal in vitro fluorescence change of chILI.1 as a function of pH. **G**) Fluorescence change, cellular brightness and affinities of main indicator variants obtained during directed evolution.

### Characterization of chILi.1

Recombinant apo-chILI.1 had two absorbance peaks at 462 nm and 562 nm, corresponding to neutral and anionic form of the cpmApple chromophore (Fig. 1B). The excitation and emission peaks were at 560 nm and 593 nm, respectively (Fig. 1C). The affinity for cAMP was ∼600 nM (n = 0.87±0.05) (Fig. 1D). The affinity for cGMP could not be accurately determined due to the lack of fluorescence change upon cGMP application, demonstrating the extraordinary specificity for cAMP over cGMP (Fig. 1D). chILI.1 had a pKa of 6.1, a very beneficial property as it facilitates stable fluorescence output in physiological pH regimes (Fig. 1E, F). In HEK293T cells chILIs expressed well, with no signs of aggregates. Expression appeared to be present in the whole cell, but higher in the nucleus. Therefore, for in-house validation and in vivo experiments, we added a C-terminal nuclear export signal and efficiently excluded fluorescence from the nucleus (Fig. 2A). For quantitative benchmarking using confocal microscopy we compared chILIs to the established red fluorescent sensors Pink Flamindo^15^ and R-FlincA^16^, imaged under identical conditions (Fig. 2B). We validated the indicators’ responses to cAMP-stimulation using 1 µM norepinephrine (NE) and 50 µM forskolin combined with 50 µM iBMX (F+I, Fig. 2A, B). Besides intensity change, chILI.1 responded to cAMP-binding with a change in fluorescence lifetime, of about 0.25 ns after norepinephrine and 0.35 ns after forskolin/iBMX (Fig. 2B). To generate a useful control sensor non-responsive to cAMP, we introduced the point mutation R91E in the cAMP-binding pocket, an amino acid exchange that would abolish binding due to drastic electrostatic charge reversal^23^. Introduction of R91E into chILI.1 successfully abolished fluorescence intensity and lifetime responses to cAMP (Fig. 2B). Finally, we assessed cellular brightness of cHILIs in HEK293T using confocal microscopy and flow cytometry. We created mTagBFP2-referenced constructs of the fluorescent proteins mApple^21^ and cpmApple, and the respective cAMP indicators (Fig. 2C). In cAMP-bound state, cHILI.1 reached 42% of mApple brightness (44% with confocal microscopy), while cpmApple, Pink Flamindo, R-FlincA and chILI.02 reached 5% (4%), 3% (3%), 8% (9%) and 9% (20%) respectively.

**Figure 2:**
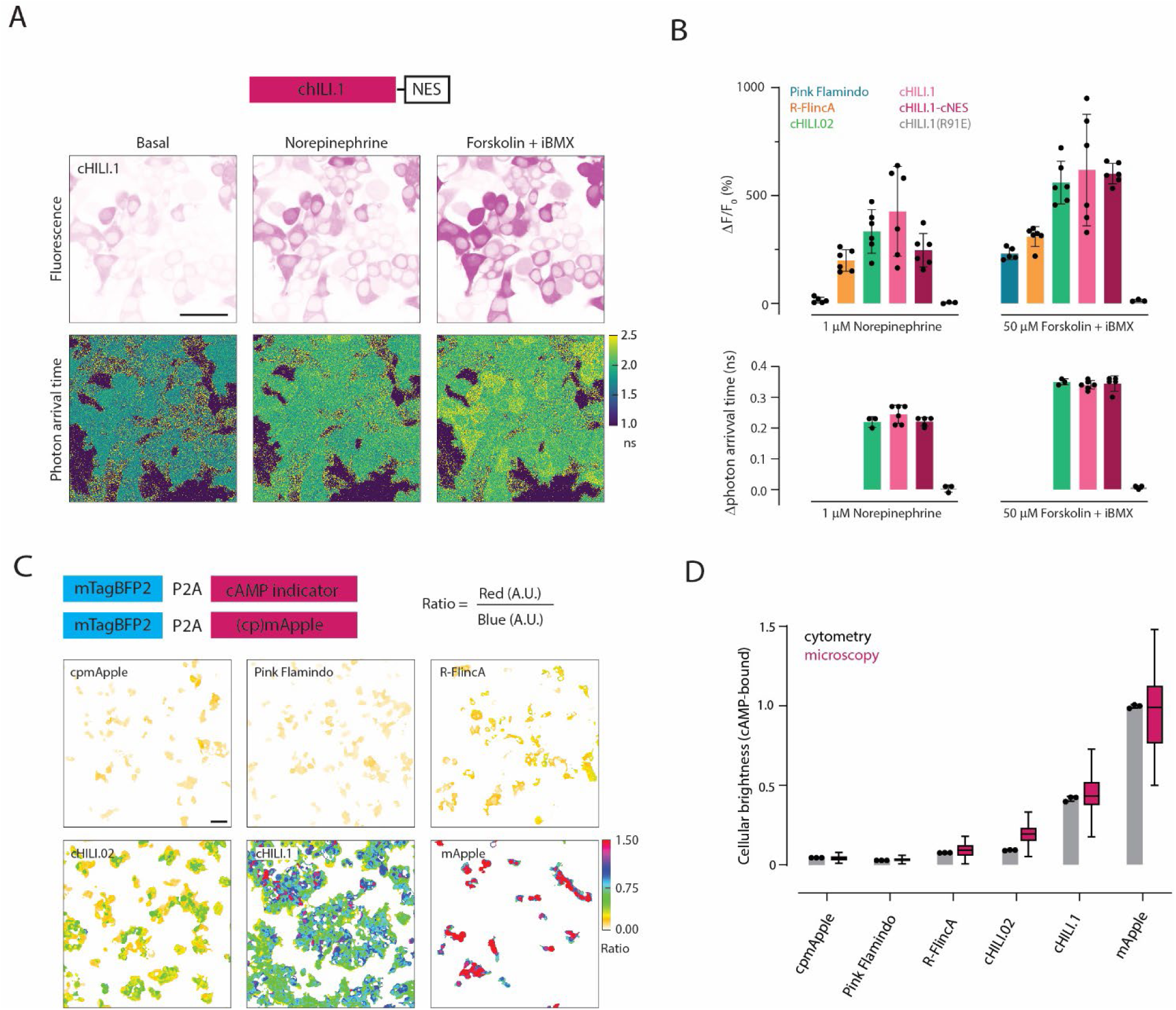
Validation of cHILI in mammalian cells. **A)** For mammalian cell validation, a nuclear export signal (NES) was added to the C-terminus of cHILI.1 (top). Representative images of cHILI.1-cNES in HEK293 cells under basal (left) and stimulated conditions, with 1 µM norepinephrine (middle), or 50 µM forskolin and 50 µM iBMX (right). Fluorescence (upper) and photon arrival time (bottom) channels are shown. Scale bar, 50 µm. **B)**. cHILIs were benchmarked against Pink Flamindo and R-FlincA. Peak responses to norepinephrine and forskolin+iBMX stimulations as normalized fluorescence change (top) or change of photon arrival time (bottom). cHILI.1(R91E), incapable of binding cAMP, was used as negative control. Dots are median responses of experimental replicates, summarized in a bar plot as mean ± SD. **C)** Scheme of referenced constructs generated by fusing fluorescent proteins and indicators to mTagBFP2 (top). Red-blue ratio was used as a proxy for cellular brightness (bottom). Representative ratioed confocal images of fluorescent proteins and indicators in the cAMP-bound state. Scale bar, 50 µm. **D)** Cellular brightness of fluorescent proteins and cAMP-indicators measured via cytometry (bars) and confocal microscopy (boxes). Bars are mean ± SD of median values obtained with experimental replicates (dots). Boxes are showing IQR with line at median, and whiskers extending to 1.5 × IQR on each side. Data is from one experimental replicate (images from C).

### Two-photon lifetime imaging of chILI.1

To characterize the relationship between fluorescence intensity and lifetime for the cpmApple-based sensor and to see cAMP dynamics in neurons using chILI.1 in advance of in vivo imaging, we next imaged the sensor activity in HEK293 cells and hippocampal pyramidal neurons with two-photon fluorescence lifetime imaging microscopy (2pFLIM) (Fig. 3). Similar to Figures 2A and 2B, chILI.1 expressed in HEK cells exhibited significant changes in fluorescence intensity and lifetime in response to cAMP stimulation (Fig. 3A, B). The relationship between lifetime and intensity change was rectangular hyperbolic, not linear (Fig. 3E, Fig. S3). Depending on the range of cAMP concentration, one can choose the parameter that is more favorable for measurement: lifetime or intensity. The sensor in the apical dendrite of hippocampal neuron also responded to cAMP stimulation using norepinephrine and forskolin (Fig. 3C, D). The basal lifetime in neurons ranged from 2.02 to 2.06 ns, identical to the 2.04 ns observed in HEK cells. Furthermore, the plot of lifetime vs intensity change exhibited a hyperbolic trend, consistent with HEK cells, except for a decrease in lifetime at the end, which was observed in all cases (Fig 3E).

**Figure 3:**
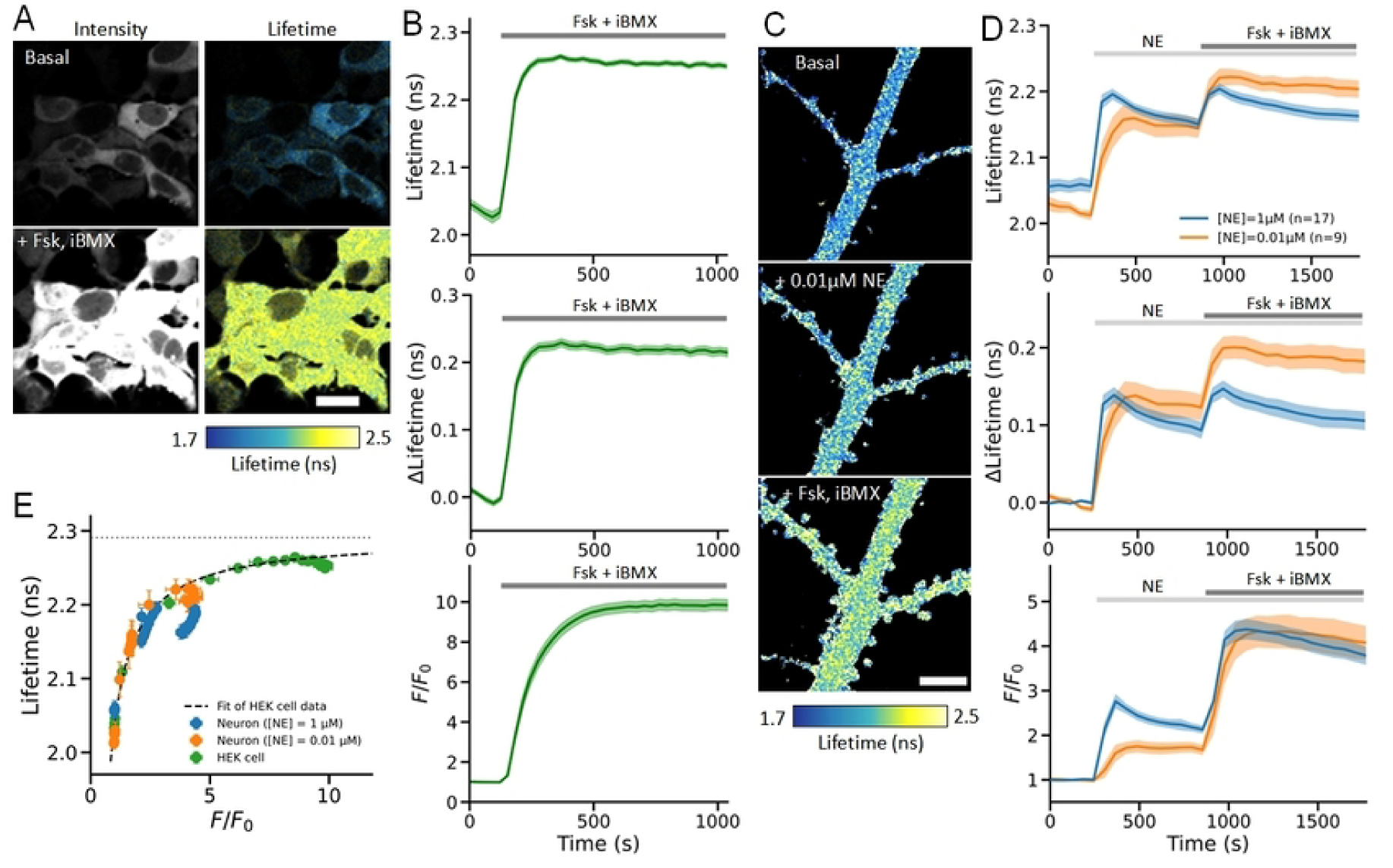
Two photon lifetime imaging of chILI.1 in HEK cells and hippocampal neurons. **A)** Representative fluorescence intensity and lifetime images of HEK293 cells expressing chILI.1 before and after forskolin stimulation. Scale bar, 20 µm. **B)** Averaged time courses of fluorescence lifetime (top), lifetime change (middle), and intensity change (bottom) in HEK cells (n = 68) treated with 50 µM forskolin containing 50 µM iBMX. **C)** Representative fluorescence lifetime images of apical dendrite in hippocampal neuron during the applications of 0.01 µM norepinephrine, followed by 25 µM forskolin containing 50 µM iBMX. Scale bar, 5 µm. **D)** Averaged time courses of fluorescence lifetime (top), lifetime change (middle), and intensity change (bottom) in neurons treated with 1 or 0.01 μM norepinephrine (n = 17 or 9 cells), followed by 50 µM forskolin containing 50 µM iBMX. **E)** Relationship between fluorescence lifetime and intensity (replotted data from B and D). Dashed line indicates a fit curve obtained from data in HEK cells (Fig.S3). Horizontal dotted line shows an extrapolated lifetime, 2.29 ns. Shaded areas in time courses and error bars in E indicate s.e.m..

### chILI. 1 performance in hippocampal CA1 of awake mice

To test the indicator’s functionality in vivo, we expressed it in mouse hippocampal CA1 via viral (AAV) injection. Mice were subsequently implanted with a chronic hippocampal window and the expressing neurons were imaged using two-photon microscopy while the head-fixed mice ran on a treadmill in the dark^24, 25^ (Fig. 4A-B). To assess the spatial extent of cAMP signaling across the field of view (FoV), we divided the FoV into 16 rectangular regions of interest (ROIs; Fig.4C) and generated ΔF/F vs time traces for each ROI. Under these conditions, we observed spontaneous cAMP transients lasting tens of seconds (Fig. 4D). In the example recording shown in Fig 4D, transients were detected in 6 of the 16 ROIs, all located in the medial portion of the FoV, whereas no detectable events were observed in the remaining ROIs (Fig. 4D).

**Figure 4:**
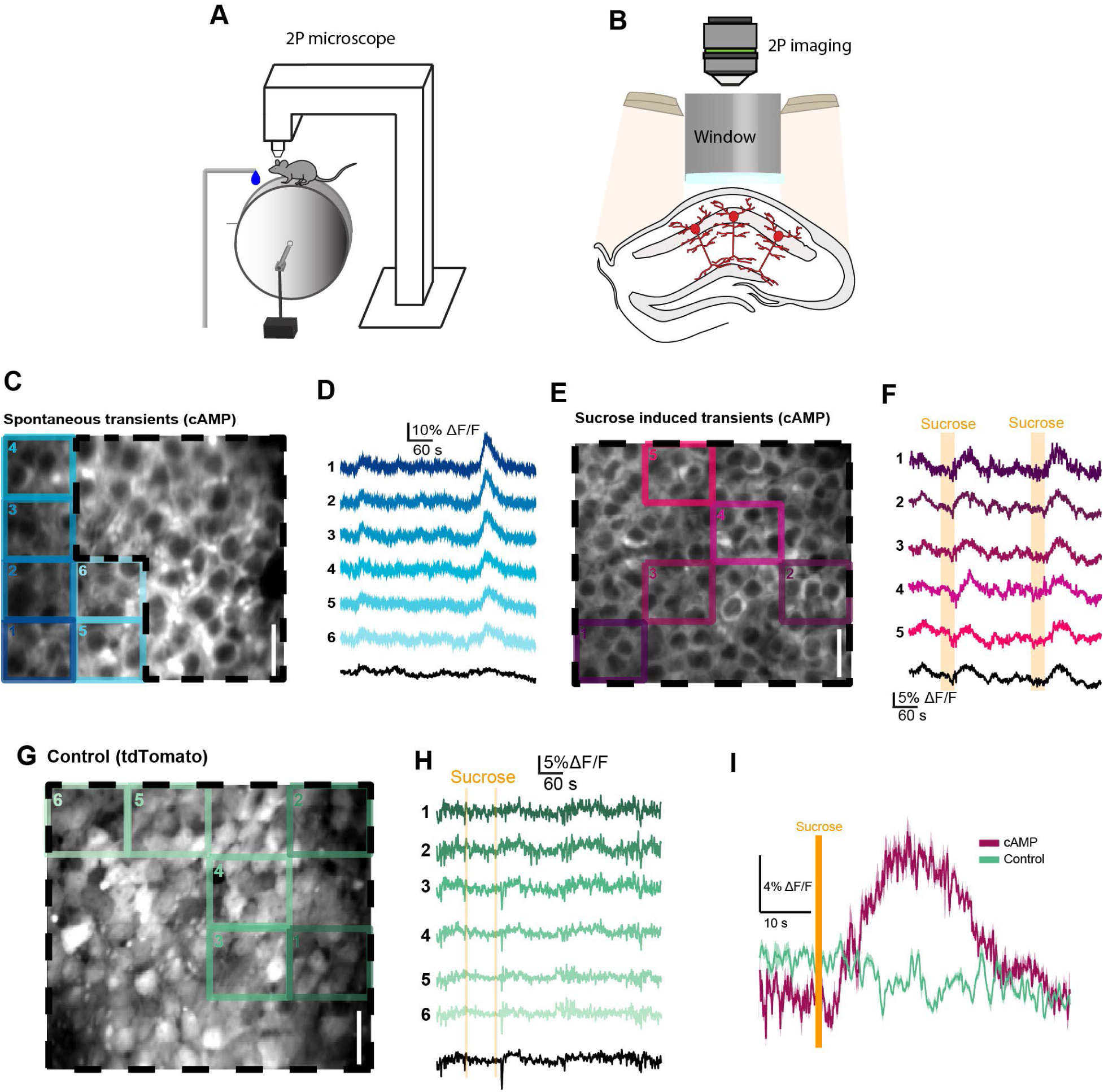
chILI.1 transients in hippocampus of awake mice. **A**) Diagram of the experimental set-up, consisting of a head-fixed mouse running on a cylindrical treadmill, and receiving sucrose-water during two photon (2P) imaging. **B**) Imaging of dorsal CA1 hippocampal neurons through a hippocampal window. **C**) Mean image of a CA1 field of view from a mouse expressing chILI.1 after AAV-mediated gene transfer with example regions of interest (ROIs). Scale bar, 30 µm. **D**) cHILI.1 transients from the ROIs shown in C. Trace colors correspond to ROI colors; the black trace corresponds to the ROI outlined with a dashed frame. **E**) Example ROIs from another mouse expressing chILI.1. Scale bar, 30 µm. **F**) chILI.1 traces from the ROIS shown in E. trace colors correspond to ROI colors; the black trace corresponds to the full-field-of-view ROI outlined with a dashed frame. Yellow rectangles indicate sucrose delivery windows. **G**) Mean image of a CA1 field of view form a mouse expressing tdTomato, with example ROIs. Scale bar, 30 µm **H**) tdTomato control traces from the ROIs shown in G. **I**) Event-triggered averages aligned to the end of the sucrose delivery window for chILI.1

We next asked whether cAMP transients could be evoked. We chose to use sucrose delivery via the drinking water. Sucrose water was delivered directly to the mouse’s mouth, and its consumption was followed by widespread cAMP transients across the FoV (Fig. 4E, F). These evoked transients were similar in duration to the spontaneous transients (Fig. 4D, F). As a control, we repeated this experiment in a mouse expressing tdTomato (a fluorescent protein inert to cAMP) and observed no comparable (spontaneous or evoked) fluorescence transients (Fig. 4G, H). To quantify this difference, we calculated event-triggered averages of the cAMP and tdTomato control ΔF/F aligned to sucrose consumption (Fig. 4I). The cAMP ΔF/F showed a clear increase following sucrose consumption, whereas no corresponding increase was observed in the tdTomato control.

## Discussion

Here we present chILI, a red fluorescent sensor for cAMP. It was optimized via iterative rounds of directed evolution in microbial and mammalian expression and selection systems. There are several beneficial features in our view. One may be the high specificity for cAMP over cGMP, to our knowledge among the highest among current cAMP sensors. Furthermore, the single ligand binding site abolishes co-operativity while also reducing buffering of cAMP transients within cells. Finally, the relatively high brightness of the bound state is helpful for increasing photon output when recording signals under in vivo imaging conditions. Like with other sensors based on the mApple^20^ chromophore, multiplexing with other green or cyan/yellow sensors is not a problem given the appropriate hardware configuration is available. chILIs also show a distinct change in fluorescence lifetime when binding ligand, a property that offers options to use the quantitative nature of fluorescence lifetime imaging. The relationship between lifetime and intensity change was not linear, but hyperbolic. Depending on the range of cAMP concentration, one may choose the parameter that is more favorable for measurement. Overall, chILIs will provide useful options for deciphering the intricacies of the spatiotemporal dynamics of cAMP in living cells and organisms.

## Materials and Methods

### Cell lines

HEK293T (ATCC CRL-3216) cells were grown in normal growth media, Dulbecco’s Modified Eagle Medium with high glucose, pyruvate (DMEM, Gibco) supplemented with 10% fetal bovine serum (FBS, Gibco), 10 000 units penicillin, 10 000 µg streptomycin, and 29.2 mg/mL L-glutamine in 0.85% saline (PSG, Gibco) at 37°C with 5% CO_2_. HEK293FRT Cas9 cell line^22^ growth medium was additionally supplemented with 250 µg/mL G418 (Thermo Fisher).

### Directed evolution in mammalian cells

#### Generation and diversification of cell lines

Generation of the cell line with a frame-shifted target protein was performed as previously described^22^. Briefly, HEK293FRT Cas9 cells were co-transfected with plasmids encoding for Flp-recombinase and frame-shifted target protein in 1:9 mass ratio using Lipofectamine 3000 (Invitrogen) according to the manufacturer’s instructions. From the third day post transfection, cells were grown in selection media with 50 µg/mL Hygromycin B (Thermo Fisher).

For library generation, 16 culture plates (5×10^6^ cells / plate) were co-transfected each with 10 µg single guide (sg)RNA and 10 µg single-stranded oligoDNA (ssODN) using Lipofectamine MessengerMax (Invitrogen) according to the manufacturer’s instructions. sgRNA and ssODN (Metabion) were designed and prepared as in reference 22. Cells were pooled next day and FACS-sorted 72 h after transfection.

### Fluorescence activated cell sorting (FACS)

Cells were collected with trypsin (Gibco) and diluted in cold Hank Balanced Salt Solution (HBSS, Gibco) with 1 mM MgCl_2_ (Sigma). When sorted in high cAMP conditions, cells were treated with 50 µM Forskolin (Sigma or MP Biomedicals) and 50 µM 3-isobutyl-1-methylxanthine (iBMX, Sigma). Cell sorting was performed on a FACSAriaIII (BD Biosciences), operated with the software FACSDiva v6.1.3. Forward-(FSC) and side-scatter (SSC) were measured with 488 nm laser and 488/10 bandpass filter. mTagBFP2 was measured with 405 nm laser for excitation and a 450/50 bandpass filter for emission (PMT gain 310). cHILI was measured with 561 nm laser for excitation and a 582/12 bandpass filter for emission (PMT gain 494). Negative control (frame-shifted cell line before library generation) was measured every time to select the single cell population and the intensity gates. The following gating strategy was used: (1) CELLS were separated from debris by FSC-A and SSC-A, (2) SINGLE CELLS were identified by the ratio FSC-H vs. FSC-A, (3) target population (Fig. S1) was sorted in tube using a 100 µm nozzle and precision set to YIELD. In the last FACS round, selected single cells were sorted in 96-well plates. In each round, 1×10^6^ cells were recorded for population analysis in FlowJo v10.8.1 (FlowJo, LLC). Cells were collected in DMEM+PSG with 20% FBS and recovered for 24 h before switching to selection media.

### Microscopy

Final selected cell population was imaged in 96-well plates (Greiner) on a PerkinElmer Opera Phenix high-content screening system equipped with 2 sCMOS 4.4 MP cameras (Oxford Instruments / Andor Zyla 5.5 CL3) using a 20x (0.4 NA) objective, operated with Harmony software v5.2. Cells were imaged in HBSS with 1 mM MgCl_2_ and DRAQ5 (2.5 µM, Thermo Fisher) nuclear stain. Image acquisition was performed in non-confocal mode as follows: mTagBFP2 (Ex375/Em435-480), cpmApple (Ex561/Em570-630) and DRAQ5 (Ex640/Em650-760). Lasers were used at maximum power with 500-800 ms exposure time. Cells were imaged before and after addition of 50 µM Forskolin and 50 µM iBMX. For analysis, cells were identified based on cpmApple signal, objects filtered by size (250-800 µm^2^) and roundness (>0.3). Mean cpmApple and mTagBFP2 fluorescence intensity, and cpmApple/mTagBFP2 ratio per cell were measured. Cell data was aggregated as median value per well separated by time point, and used to calculate normalized ratio change ΔR/R_0_ (%) in each well. Copy of the samples was kept in culture and gDNA was extracted from selected variants to identify the indicator gene sequence.

### Directed evolution in bacterial cells

DNA libraries were generated using error prone or site saturation PCR and cloned into pRSET-B vector, incorporated into *E. coli* XL1 Blue cells (Invitrogen) and transferred onto 20-40 LB-Agar plates supplemented with 50 µg/mL Ampicillin (Roth). Plates were incubated overnight at 37⁰C and then at 4⁰C for 48 h to favor protein folding. Plates were imaged using a home-built screening platform^26^ using 550/10 nm excitation (1-3 s exposure) and 600/10 nm emission filters. Images were acquired and analyzed using customized python routines. Colonies were selected based on the red fluorescence intensity. DNA of the selected colonies was isolated, the sensors were expressed and purified for further characterization in solution. Variants were filtered according to response to 1 mM cAMP (DF/F_0_) and extinction coefficient (EC). If a suitable candidate was identified, its response was also evaluated in HEK293T cells.

### Protein purification and biophysical characterization

His-tagged proteins were expressed in *E. coli* BL21 (Invitrogen) overnight at 37°C in 50 mL auto-inductive Luria-Bertani medium, supplemented with 0.05% D-(+)-glucose (w/v), 0.2% lactose (w/v), 0.6% glycerol (v/v). Bacterial cells were harvested by centrifugation (4°C, 10 min, 6 000 *g*) and resuspended in 10 mL resuspension buffer (20 mM Na_2_PO_4_, 300 mM NaCl, 20 mM imidazole; Sigma Aldrich) supplemented with protease inhibitors (4 μM PMSF, 20 μg/mL Pepstatin A, 4 μg/mL Leupeptin; Sigma Aldrich), 5 μg/mL Dnase, and 10 μg/mL RNase (Sigma Aldrich). After resuspension, bacteria were lysed by sonication on ice for 7 min (0.8 cycle, 80% power; Bandelin Sonoplus). After centrifugation (4°C, 30 min, 20 000 *g*), the supernatant was incubated with 140 μL 6% (v/v) Nickel-IDA agarose bead suspension (Jena Bioscience) for 60 min at 4°C under mild agitation. Agarose beads were collected using 1 mL propylene gravity flow columns (Qiagen) and washed twice with 5 mL resuspension buffer. Finally, proteins were collected using 600 μL elution buffer (20 mM Na_2_PO_4_, 300 mM NaCl, 300 mM imidazole; Sigma Aldrich). Experimental triplicates (protein productions) were done for each indicator and fluorescent protein.

The fluorescence change (ΔF/F_0_ %) of purified cHILI variants was determined by measuring the fluorescence at 610 nm upon excitation at 570 nm in HEPES buffer (25 mM HEPES, 200 mM NaCl, pH 7.5) supplemented with 1 mM cAMP and 1 mM MgCl_2_. The molar extinction coefficient (EC) was determined using the alkaline protocol, in which the protein concentration is obtained using the absorption of the denatured chromophore at 452 nm (EC of 44 mM^−1^cm^−1^). Proteins were prepared in HEPES buffer supplemented with 1 mM cAMP and the absorbance spectrum was acquired before and after addition of NaOH to a final concentration of 0.1 M. The quantum yield was determined relative to mApple using the slope method, measuring the absorbance and emission spectra of serial dilution of the proteins. The brightness was calculated as extinction coefficient ^ˣ^ quantum yield.

The cAMP/cGMP (cNMP) binding affinity of the sensors was determined using HEPES buffer supplemented with 1 mM MgCl_2_ by serially decreasing concentrations of cNMP 1:4, starting from 1 mM. Briefly, 0.5-1 μM protein was prepared in buffer with 1 mM or no cNMP. 200 μL of protein with 1 mM cNMP was pipetted in the first well of a 96-well plate, and 150 μL of protein with no cNMP was added to the subsequent wells. 50 μL of high cNMP mixture was transferred to the neighboring well and resuspended, resulting in 1:4 dilution of cNMP. This step was repeated until 11 serial dilutions were made, covering the range from 1 mM to 0.95 nM cNMP. The dissociation constant (Kd) values were determined by plotting the log10 values of the cNMP concentrations against the corresponding ΔF/F_0_ values (normalized to the max response) and fitting a sigmoidal curve to the plot.

For measuring the pKa of the sensors a series of MOPS/TRIS/Citrate buffered solutions, supplemented with either 1 mM or no cAMP, were prepared. The pH values were adjusted in 0.5 pH steps from pH 4.5 to pH 8.5 using NaOH and HCl. In a 96 well plate, 200 μL of buffer with 0.5-1 μM of protein were prepared for each pH value and emission spectra were recorded. The relative fluorescence values at the emission maximum were plotted against the pH values and a sigmoidal fit was applied.

### Evaluation of cAMP responses in HEK293T cells

To determine fluorescence response, cells were seeded in 35 mm glass-bottom dishes (Mat-Tek) pre-coated with poly-L-lysine (Sigma) or on 96-well plates (Ibidi) and transfected with Lipofectamine 3000 one day before imaging. Time-lapsed images were acquired on Leica SP8 Matrix or Leica Stellaris5 microscope (both Leica) with 20x water-dipping (NA1.00, SP8), oil-immersion (NA0.75, Stellaris5) or air objective (NA0.75, Stellaris5), with excitation at 561 nm and emission in the range 580-650 nm. Cells were imaged in HBSS supplemented with 1 mM MgCl_2_. Responses to bath application of 1 mM norepinephrine and 50 mM Forskolin + 50 mM iBMX (final concentrations) were measured. Arrival time data was acquired on Leica Stellaris5, using TauContrast mode of the HyDS2 detector with 4 line accumulations to increase the photon count per pixel. Data was acquired from at least three experimental triplicates (transfections).

### Evaluation of indicator brightness in HEK293T cells

To determine the molecular brightness in mammalian cells, mTagBFP2 was fused at the N-terminus of selected variants or fluorescent proteins, separated by a P2A peptide. Indicators were compared to the mTagBFP2-P2A-mApple construct, using confocal microscopy (Leica Stellaris5, Leica) and flow cytometry (Attune NxT Analyzer, Thermo Fisher). Brightness was obtained in the presence of 50 mM Forskolin + 50 mM iBMX.

Microscopy was done as described for the fluorescence response, but acquiring two channels: blue channel (mTagBFP2) with excitation at 405 nm and emission between 440-480 nm, and red channel as described above. The ratio F_Ex561_/F_Ex405_ was calculated for each variant. Data was acquired from one experimental replicate (transfection).

For flow cytometry, HEK293T cells were seeded in 6-well plates (Greiner) and transfected with the different mTagBFP2-P2A-constructs using Lipofectamine 3000 24 h after seeding. Next day, cells were prepared for flow cytometry as described for FACS. All samples were recorded before and after addition of 50 mM Forskolin and 50 mM iBMX. At least 19 000 events were recorded for each condition and further analyzed using FlowJo v10.8.1 (FlowJo, LLC). Events were first gated by forward and side scattering for selecting live single cells. Attune VL1-A channel (Excitation 405 nm - Emission 440/50 nm) was used for mTagBFP2, and YL2-A (Excitation 561 nm - Emission 620/15 nm) for indicators and (cp)mApple. Ratio values (F_YL2-A_/F_VL1-A_) were obtained for each cell and normalized to the median value of the mApple group. Data was acquired from three experimental triplicates (transfections).

### Image processing and data analysis

Raw images were analyzed with Fiji v1.54p. Threshold mask was created based on red channel data in the first acquired frame to assign N/A values to the background and saturated pixels. Original images were divided by the threshold mask and a median filter with 2-pixel diameter was applied. Arrival time channel pixel values were multiplied by 0.097 and subtracted by 1 after thresholding. First frame of the red channel was segmented with cellpose3 v3.1.1.2^27^ using built-in cyto3 model. ROIs were saved in ImageJ supported format and overlaid onto processed images. Mean pixel intensity values were measured for each ROI across different timepoints. Data was analyzed in Excel and plotted with GraphPad Prism v11.0.0 (GraphPad Software).

### Organotypic hippocampal slice cultures

Organotypic cultured hippocampal slices were prepared from postnatal 7-8 days C57BL/6 mice, as described^28, 29^. After 7 days in culture, hippocampal pyramidal neurons were transfected via biolistic gene transfer (Bio-Rad) with gene bullets made from 1-μm gold beads (8 mg) coated with plasmid of pCAGIG-chILI.1-cNES (30 μg). 2pFLIM imaging was performed 3-6 days after transfection.

### Surgery

All mouse procedures were approved by the Northwestern University Institutional Animal Care and Use Committee. Male C57BL/6 mice were singly housed under a reverse light–dark cycle, with experiments performed during the dark phase, at 20–24 °C and 30%–70% humidity. At 9 weeks of age, mice were anesthetized with 1%–2% isoflurane. A 1-mm diameter craniotomy was made over the right hippocampus, centered 1.8 mm lateral and 2.3 mm caudal to bregma. A beveled glass micropipette was lowered to a depth of 1.25 mm below the brain surface, and 80 nl of adeno-associated virus was injected at each of two sites located 150 µm anterior and posterior to the center of the craniotomy. The cAMP sensor was expressed using cNES-pAAV-hSyn1-WPRE-pA, whereas control mice were injected with pAAV-CAG-tdTomato. Viral titers were 7.36 × 1012 and 5.75 × 1012 genome copies per ml for cAMP sensor and tdTomato control, respectively.

The mice then underwent water scheduling, receiving 0.8–1 ml of water per day for the duration of the study. Nine days after viral injection, a chronic hippocampal imaging window was implanted as described previously^25^.

### Two-photon fluorescence lifetime imaging

Fluorescence lifetimes were measured on a custom-built 2pFLIM system^29, 30^ equipped with a Ti:Sapphire laser (Spectra-Physics, InSightX3) with a fixed wavelength of 1040 nm. The laser power was modulated using a Pockels cell to 3.3-4.1 mW under the objective lens, and the laser position was scanned using XY galvo with 3-mm mirrors (Cambridge Technology, Model 6215H). Experiments were conducted in a HEPES-buffered ACF (20 mM HEPES pH 7.3, 130 mM NaCl, 2 mM NaHCO_3_, 25 mM D-glucose, 2.5 mM KCl and 1.25 mM NaH_2_PO_4_) with 2 mM CaCl_2_ and 2 mM MgCl_2_. Neurons were stimulated with bath application of 1 or 0.01 μM norepinephrine (Tocris), followed by 25 µM forskolin (Tocris) combined with 50 µM iBMX (Tocris). The red fluorescence was collected with a water-immersion objective (Olympus, LUMPlanFLN60× NA 1.0 W), separated by a dichroic mirror (565 nm) and detected with a photoelectron multiplier tube (Hamamatsu Photonics, H7422-P40) placed after emission filters (Chroma 620/60-2p). Fluorescence lifetime image was obtained using a time-correlated single-photon counting board (Pico-Quant, Time-Harp 260) controlled with custom software (FLIMage)^29^. 2pFLIM images of apical dendrite were acquired at 1-minute intervals for a 23.2×23.8 μm area with a resolution of 256×256 pixels, an average of 12 frames, and a Z-stack of five slices (1 µm pitch). Experiments were performed at room temperature (23 ± 1 °C).

2pFLIM images of HEK293 cells were taken of a 232.0×237.5 μm area at 256×256 pixels with a 24-frame average at 30-second intervals. The cells were stimulated with bath application of 50 µM forskolin containing 50 µM iBMX. ROIs for cells were generated by Cellpose-SAM^31^ for lifetime analysis.

### Two-photon in vivo imaging

For in vivo imaging, mice were head-fixed on a cylindrical treadmill and maintained in darkness. High-volume white noise was played in the experimental room to minimize external sensory stimuli. Sucrose water was delivered manually to the mouth of the head-fixed mouse using a transfer pipette. Imaging was performed using a custom movable-objective microscope (Sutter Instruments) equipped with a resonant scanning module (Sutter Instruments) and a 40×/0.8 NA water-immersion objective (LUCPlanFL N, Olympus). Excitation was provided by a fixed-wavelength 1064-nm laser (AXON 1064-3 TPC, Coherent), and imaging was controlled using ScanImage software (Vidrio Technologies). Time-series movies were acquired using bidirectional scanning at 60 Hz with an image size of 512 × 256 pixels.

### ROI definition and fluorescence trace extraction

In vivo movies were motion-corrected using whole-frame cross-correlation, as described previously^25, 32^. Each imaging field of view was divided into a 4 × 4 grid of rectangular regions of interest (ROIs), and the mean fluorescence signal was calculated for each ROI. Fluorescence traces were converted to ΔF/F using an 8th-percentile baseline calculated within a 150-s sliding window^24^.

### Analysis

Event-triggered averages of ΔF/F traces were calculated by aligning fluorescence signals to the times of sucrose-water delivery. This analysis was performed separately for the cAMP indicator and tdTomato control recordings. For each condition, the event-triggered average was calculated across 16 ROI traces and two sucrose-delivery events.

## Acknowledgements

This work was supported by NIH Grant UO1 NS128655-01 (to D.D., R.Y. and O.G.) and by the Max-Planck-Society. We want to acknowledge the following service facilities of the MPI for Biochemistry: The Biochemistry Core Facility (RRID:SCR_025743) and the Imaging Facility (RRID:SCR_025739). We also acknowledge the imaging facility of the MPI for Biological Intelligence (RRID:SCR_026797) for service and equipment use. Furthermore, we would like to thank Anja Moritz for technical assistance.

---

**Supplementary Figure S1.**
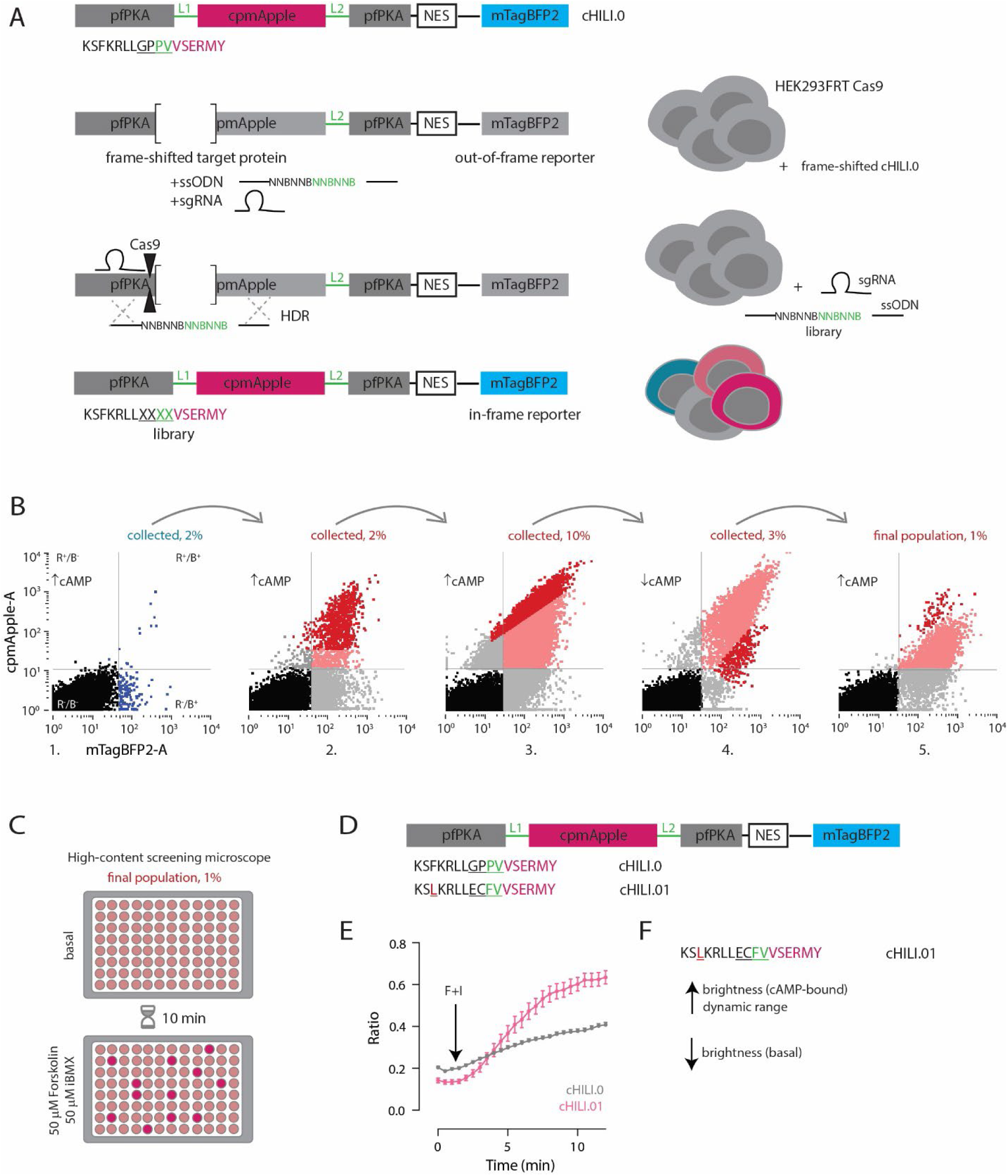
Directed evolution of linker 1 region in mammalian cells. **A)** Library generation in mammalian cells. cHILI.0 prototype was C-terminally fused to mTagBFP2, a reporter of library cells. Deletion causing a shift of reading-frame was made near the linker 1 region, and resulting dark construct was integrated into the HEK293FRT Cas9 cell line. Library was generated by HDR after a double-strand DNA break. Cells were provided with sgRNA targeted near the frame-shift site and single strand DNA oligo library with overhangs complementary to the region around the break. HDR resulted in repair of reading frame and recovery of fluorescence. **B)** Strategy for directed evolution of linker 1 during five consecutive FACS rounds. Quadrants on each scatterplot display four different red-blue subpopulations determined by the negative control. Dark pink or blue, light pink, gray and black dots represent collected, discarded double-positive, discarded single-positive, and double-negative cells, respectively. Arc arrows show the direction of the screening rounds. In rounds 1, 2, 3, and 5 cells were stimulated with 50 µM forskolin and 50 µM iBMX. Final population was collected as single cells in two 96-well plates. **C)** Screening of the final sorted population was done with the Opera Phenix high-content screening microscope. After a baseline image, cells were stimulated with forskolin and iBMX, and imaged again after 10 min. **D)** Amino acid changes of the best performing variant cHILI.01, compared to cHILI.0. Note the unexpected mutation F111L. **E)** Red-blue ratio of cHILI.01 and cHILI.0 before and after stimulation with 50 µM forskolin and 50 µM iBMX. **F)** Compared to cHILI.0, cHILI.01 has lower basal and higher cAMP-bound brightness, resulting in improved dynamic range.

**Supplementary Figure S2.**
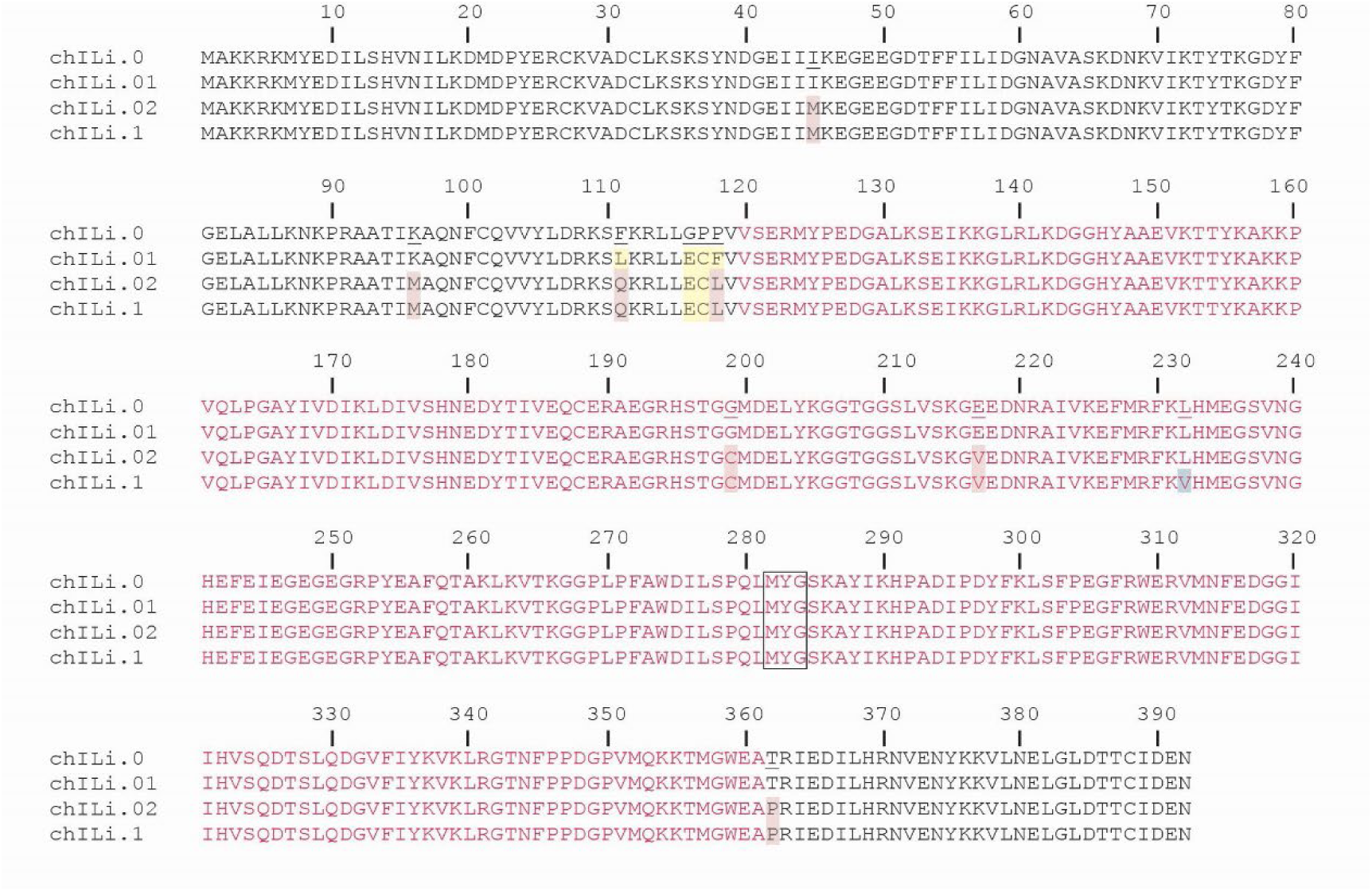
Amino acid sequences of chILI.1 and selected parental variants. pfPKA residues are in black, cpmApple residues in red. The cpmApple chromophore is boxed.

**Supplementary Figure S3.**
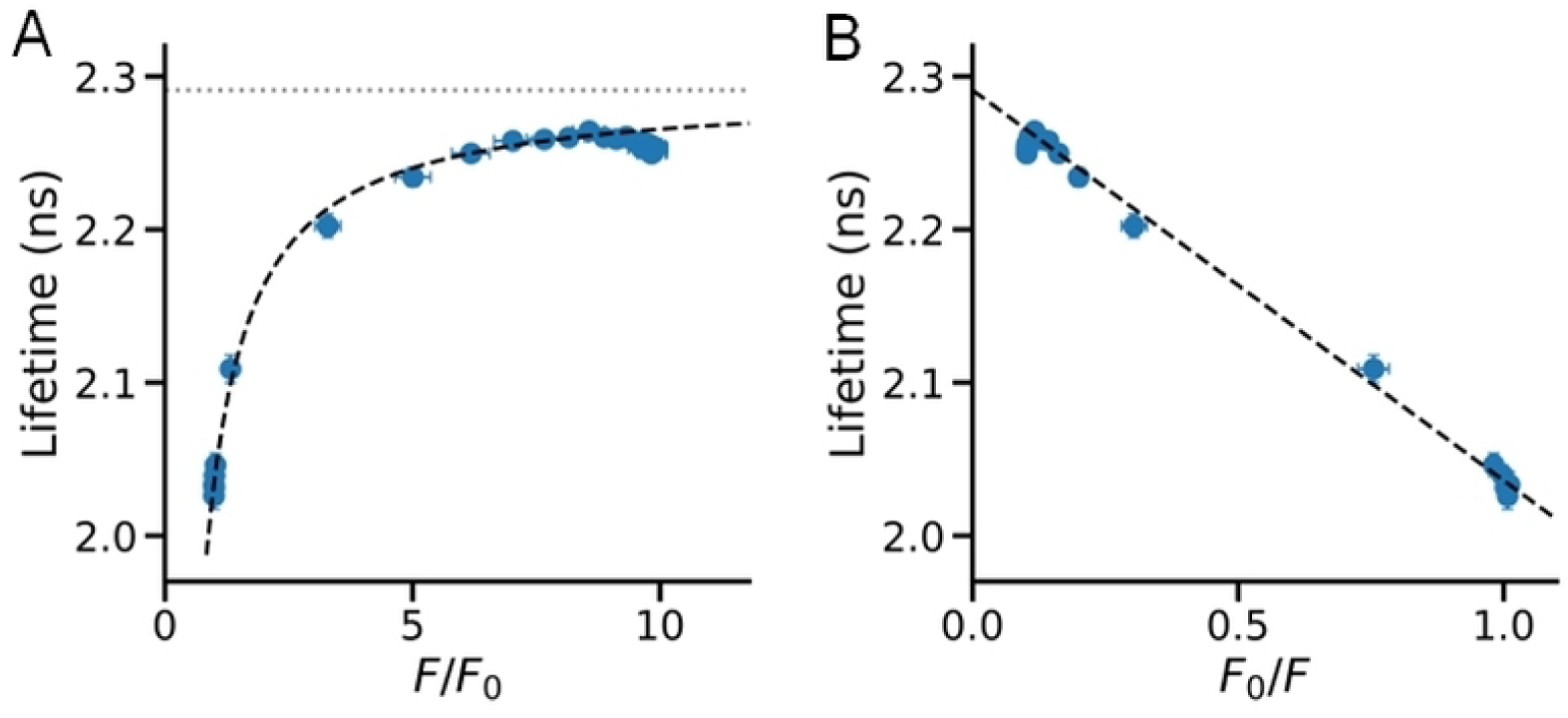
Relationship between fluorescence lifetime and intensity in a two-state model. **A)** Fluorescence lifetime vs *F*/*F*_0_ in HEK293 cells represents a rectangular hyperbola manner according to Equation (5) below. The dashed line shows a fit curve in the range of *F*/*F*_0_ < 9 (*τ*_∞_ = 2.291 ± 0.002, *τ*_base_ = 2.036 ± 0.003, *χ*^2^/dof = 0.65, R^2^ = 0.9972). **B)** The linear representation of fluorescence lifetime vs *F*_0_/*F*. The intercept *τ*_∞_ is 2.290 and the slope is 0.254. Error bars indicate s.e.m..

To derive the equation of the relationship between lifetime and intensity, we model the sensor as having two fluorescent states—a dim state A and a bright state B—with a fraction *x* of molecules in state B. Assuming a single-exponential fluorescence decay for each state, the photon arrival-time distribution is

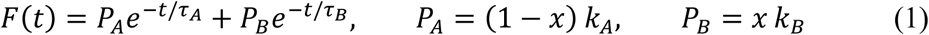

where *τ_i_* is the fluorescence lifetime of state *i* and *k_i_* is an amplitude proportional to the product of that state’s excitation cross-section and radiative rate constant. The integrated intensity and mean lifetime, defined by the mean photon arrival time (Eq. 5 in Yasuda 2006), are

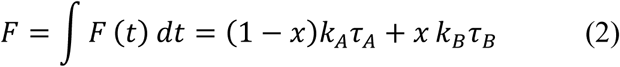

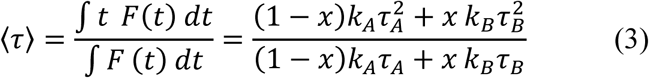

Both *F* in (2) and the numerator of (3), ‹*τ*›*F*, are linear in *x*. Eliminating *x* yields a linear relationship between these quantities:

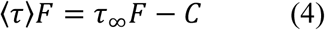

Where *τ*_∞_ = (*k_B_ τ_B_*^2^ – *k_A_ τ_A_*^2^)/(*k_B_ τ_B_* – *k_A_ τ_A_*) and *C* is a constant. Both depend only on *k_A_*, *k_B_*, *τ_A_* and *τ_B_*, and are independent of *x*.

Let *F*_0_ and *τ*_base_ denote the intensity and mean lifetime of an arbitrary reference (baseline) state, with a fraction *x*_0_ of molecules in state B. Equation (4) also holds for this reference state: C = (*τ* _∞_ − *τ*_base_) *F*_0_. Substituting this relation into (4) and dividing by *F* gives

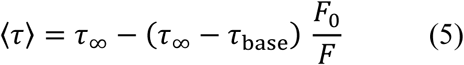

Equation (5) holds for any reference state; the baseline need not contain only molecules in state A.

Equation (5) describes ‹*τ*› as a rectangular hyperbola in *F*/*F*_0_ with asymptote *τ*_∞_ and as a straight line in *F*_0_/*F* with intercept *τ*_∞_ and slope – (*τ*_∞_ − *τ*_base_). The linear representation provides a direct test of the two-state model. *τ*_∞_ denotes the lifetime extrapolated to ∞ and can be written as *τ*_∞_ = *τ_A_* + (*τ_B_* − *τ_A_*)*r*/(*r* − 1), where the brightness ratio is *r* = *k_B_ τ_B_*/(*k_A_ τ_A_*). When *r* >> 1, *τ*_∞_ ≈ *τ_B_* and (5) reduces to ‹*τ*› ≈ *τ*_base_ + (*τ_B_* − *τ*_base_)Δ*F*/*F*. Thus, the mean lifetime is linear in Δ*F*/*F*, the fraction of detected photons originating from state B.

## Notes

### Competing Interest Statement

The authors have declared no competing interest.

